# Data-independent acquisition method for ubiquitinome analysis reveals regulation of circadian biology

**DOI:** 10.1101/2020.07.24.219055

**Authors:** Fynn M. Hansen, Maria C. Tanzer, Franziska Brüning, Isabell Bludau, Brenda A. Schulman, Maria S. Robles, Ozge Karayel, Matthias Mann

## Abstract

Protein ubiquitination is involved in virtually all cellular processes. Enrichment strategies employing antibodies targeting ubiquitin-derived diGly remnants combined with mass spectrometry (MS) have enabled investigations of ubiquitin signaling at a large scale. However, so far the power of data independent acquisition (DIA) with regards to sensitivity in single run analysis and data completeness have not yet been explored. We developed a sensitive workflow combining diGly antibody-based enrichment and optimized Orbitrap-based DIA with comprehensive spectral libraries together containing more than 90,000 diGly peptides. This approach identified 35,000 diGly peptides in single measurements of proteasome inhibitor-treated cells – double the number and quantitative accuracy of data dependent acquisition. Applied to TNF-alpha signaling, the workflow comprehensively captured known sites while adding many novel ones. A first systems-wide investigation of ubiquitination of the circadian cycle uncovered hundreds of cycling ubiquitination sites and dozens of cycling ubiquitin clusters within individual membrane protein receptors and transporters, highlighting novel connections between metabolism and circadian regulation.

## INTRODUCTION

Ubiquitination is a reversible and highly versatile post-translational modification (PTM) involved in virtually all cellular processes. A ubiquitin conjugation cascade, involving ubiquitin activating (E1), conjugating (E2) and ligating (E3) enzymes, mediates the covalent attachment of the 76 amino acid long ubiquitin molecule to a ε-amine group of a lysine residue on a substrate protein. Its removal is mediated by an enzyme family called deubiquitinating enzymes (DUB). Ubiquitin itself can be ubiquitinated N-terminally or via one of its seven lysine residues, giving rise to a plethora of chain topologies, which encode a diverse and specific set of biological functions^1,2^. Deregulation of this highly complex process has been linked to numerous diseases including neurodegenerative diseases^3,4^, autoimmunity^5,6^ and inflammatory disorders^7–9^.

Protein ubiquitination is one of the most widely studied PTMs in the field of mass spectrometry (MS)-based proteomics. However, due to low stoichiometry of ubiquitination and varying ubiquitin chain topologies, comprehensive profiling of endogenous ubiquitination is challenging and requires one or more enrichment steps prior to MS analysis^10^. Early reports to catalog ubiquitin conjugated proteins from yeast and human described various enrichment methods including the use of epitope-tagged ubiquitin or ubiquitin-associated domains (UBA)^11–13^. After trypsinization, previously ubiquitinated or NEDDylated peptides bear a signature diGly remnant that can be targeted by a specific antibody^14^. Enrichment strategies employing such antibodies have enabled identification of thousands of ubiquitination sites by MS^15–17^. A recently described antibody targets a longer remnant generated by LysC digestion to exclude ubiquitin-like modifications such as NEDD8 or ISG15^18^, however, the contribution of diGly sites derived from ubiquitin-like modifications is very low (< 6%)^15^.

The commercialization of such antibodies has accelerated MS-based ubiquitinome analysis and enabled a variety of quantitative, systems-wide studies^19–23^. However, large-scale analysis of ubiquitination events to study key signaling components remains challenging since in-depth diGly proteome coverage requires relatively large sample amounts and extensive peptide fractionation. These requirements, which largely stem from the low stoichiometry of the modification, come at the expense of throughput, robustness and quantitative accuracy.

Thus far, ubiquitinome studies have employed data-dependent acquisition (DDA) methods combined with label-free or isotope-based quantification^24^. Recently, Data Independent Acquisition (DIA) has become a compelling alternative to DDA for proteomics analysis with greater data completeness across samples^25–28^. In contrast to intensity-based precursor picking of DDA, DIA fragments all co-eluting peptide ions within predefined mass-to-charge (*m/z*) windows and acquires them simultaneously^29^. This leads to more precise and accurate quantification with fewer missing values across samples and higher identification rates over a larger dynamic range. DIA usually requires a comprehensive spectral library, from which the peptides are matched into single run MS analyses. Recently, superior performance of DIA for sensitive and reproducible MS measurements has also been demonstrated for global protein phosphorylation analysis^30^. Given the central importance of ubiquitination, we here set out to investigate the power of DIA for improving data completeness and sensitivity in a single run analysis format.

For sensitive and reproducible analysis of the ubiquitin-modified proteome, we devised a workflow combining diGly antibody-based enrichment with a DIA method tailored to the unique properties of the library peptides and to the linear quadrupole Orbitrap mass analyzer. We acquired extensive spectral libraries that all together contain more than 90,000 diGly peptides allowing us to reproducibly analyze 35,000 distinct diGly peptides in a single measurement of proteasome inhibitor-treated cells. The DIA-based diGly workflow markedly improved the number and quantitative accuracy compared to DDA. To investigate if our new method would have advantages in the exploration of biological signaling systems, we first applied it to the well-studied TNF-signaling pathway, where it retrieved known ubiquitination events and uncovered novel ones. We then extended it to the analysis of circadian post-translational dynamics, so far poorly studied globally with regards to ubiquitination. This uncovered a remarkable extent and diversity of ubiquitination events. These include closely spaced clusters with the same circadian phase, which are likely pointing to novel mechanisms. Together, our design and results establish a sensitive and accurate DIA-based workflow suitable for investigations of ubiquitin signaling at a systems-wide scale.

## RESULTS

### DIA quantification enables in-depth diGly proteome coverage in single shot experiments

To obtain a comprehensive, in-depth spectral library for efficient extraction of diGly peptides in single shot DIA analysis, we treated two human cell lines (HEK293 and U2OS) with a common proteasome inhibitor (10 μM MG132, 4 h). After extraction and digestion of proteins, we separated peptides by basic reversed-phase (bRP) chromatography into 96 fractions, which were concatenated into 8 fractions (**Methods**, **Supplementary Fig. 1a**). Here, we isolated fractions containing the highly abundant K48-linked ubiquitin-chain derived diGly peptide (K48-peptide) and processed them separately to reduce excess amounts of K48-peptides in individual pools, which compete for antibody binding sites during enrichment and interfere with the detection of co-eluting peptides (**Supplementary Fig. 1b**). We found this to be a particular issue for MG132 treatment, as blockage of the proteasome activity further increases K48-peptide abundance in these samples. The resulting 9 pooled fractions were enriched for diGly peptides, which were separately analyzed using a DDA method (PTMScan Ubiquitin Remnant Motif (K-ε-GG) Kit, CST) (**Fig. 1a** and **Supplementary Fig. 1a-b**). This identified more than 67,000 and 53,000 diGly peptides in MG132 treated HEK293 and U2OS cell lines, respectively (**Fig. 1b**). Furthermore, to fully cover diGly peptides of an unperturbed system, we also generated a third library using the same workflow but with untreated U2OS cells (used later for biological applications). This added a further 6,000 distinct diGly peptides (**Fig. 1b**). In total, we obtained 89,650 diGly sites corresponding to 93,684 unique diGly peptides, 43,338 of which were detected in at least two libraries (**Fig. 1c**, see also source data at PRIDE: PXD019854). To our knowledge, this represents the deepest diGly proteome to date. According to the PhosphositePlus database^31^, 57% of the identified diGly sites were not reported before and 7.3% of them were previously found to be acetylated or methylated, indicating that different PTMs can act on the same sites.

**Fig. 1.**
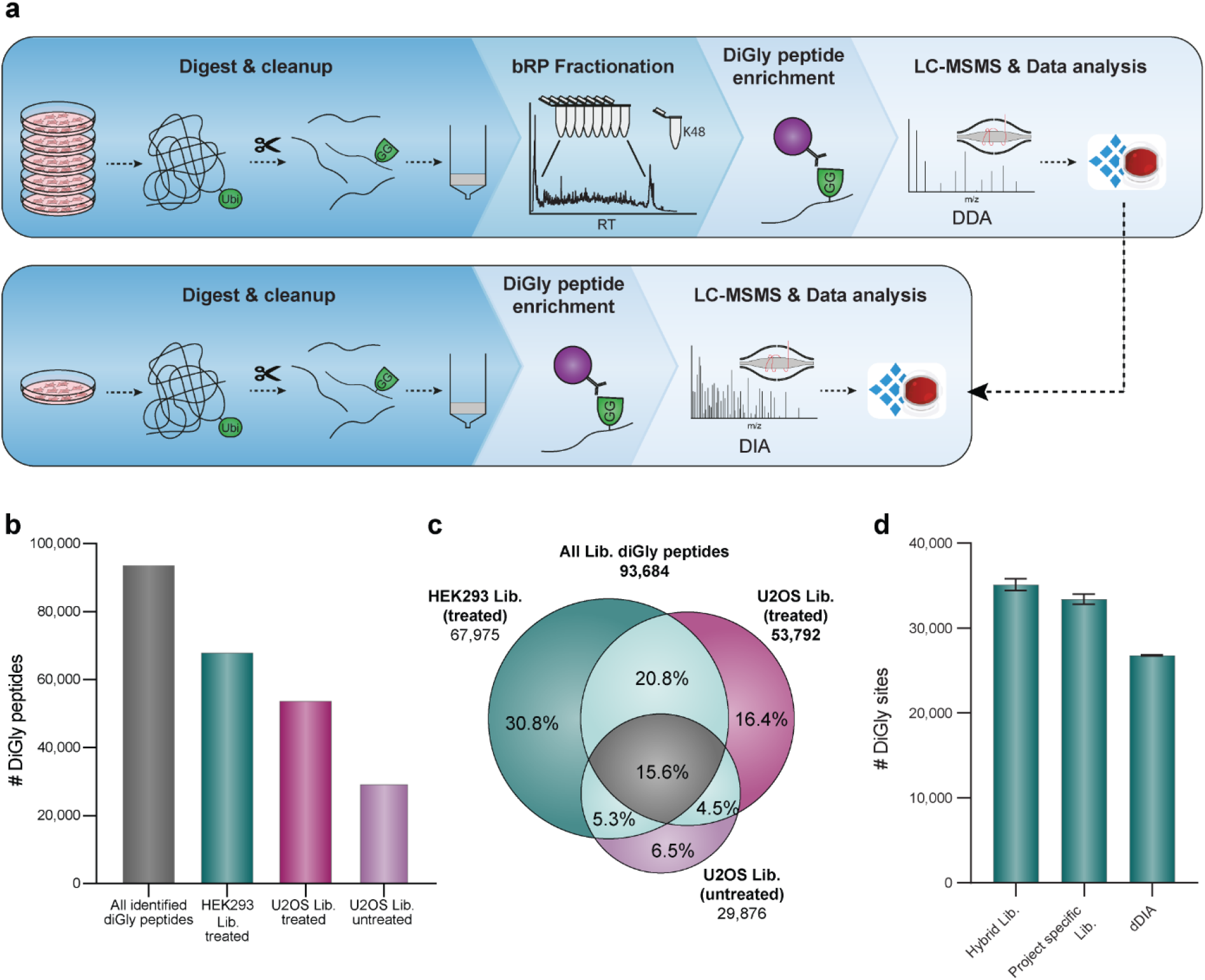
In-depth diGly proteomics for DIA identification. **a** Experimental workflow for in-depth diGly peptide library construction (upper panel) and our single run DIA-based workflow (lower panel). Protein digestion and peptide extraction are followed by bRP fractionation and diGly peptide enrichment. For library construction, samples were measured by DDA and computationally processed (Spectronaut Pulsar). Individual samples are measured by our DIA workflow, including matching against a library for identification (Spectronaut software). **b** Number of identified diGly peptides in three different spectral libraries. **c** Commonly and exclusively identified diGly peptides for different libraries. **d** Number of identified diGly sites (± SEM) of three workflow replicates of MG132 treated HEK293 cells measured in analytical duplicates using different DIA library search strategies.

In possession of these large diGly spectral libraries, we evaluated DIA method settings for best performance in single shot diGly experiments (**Supplementary Table 1**). Impeded cleavage C-terminal to modified lysine residues frequently generates longer peptides with higher charge states, resulting in diGly precursors with unique characteristics. Guided by the empirical precursor distributions, we first optimized DIA window widths – the transmission windows that together cover the desired precursor peptide range. This increased the number of identified diGly peptides by 6% (**Supplementary Fig. 2a-b**). Next, we tested different window numbers and fragment scan resolution settings, to strike an optimal balance between data quality and a cycle time that sufficiently samples eluting chromatographic peaks. We found that a method with relatively high MS2 resolution of 30,000 and 46 precursor isolation windows performed best (13% improvement compared to the standard full proteome method that we started with) (**Supplementary Fig. 2c**). Furthermore, we determined the optimal antibody and peptide input combination to maximize peptide yield and depth of coverage in single DIA experiments. To mimic endogenous cellular levels, we used peptide input from cells not treated with MG132. From titration experiments, enrichment from 1 mg of peptide material using 1/8^th^ of an anti-diGly antibody vial (31.25 μg) turned out to be optimal (**Methods** and **Supplementary Fig. 2d,e**). With the improved sensitivity by DIA, only 25% of the total enriched material needed to be injected (**Supplementary Fig. 2f**).

**Table 1.**
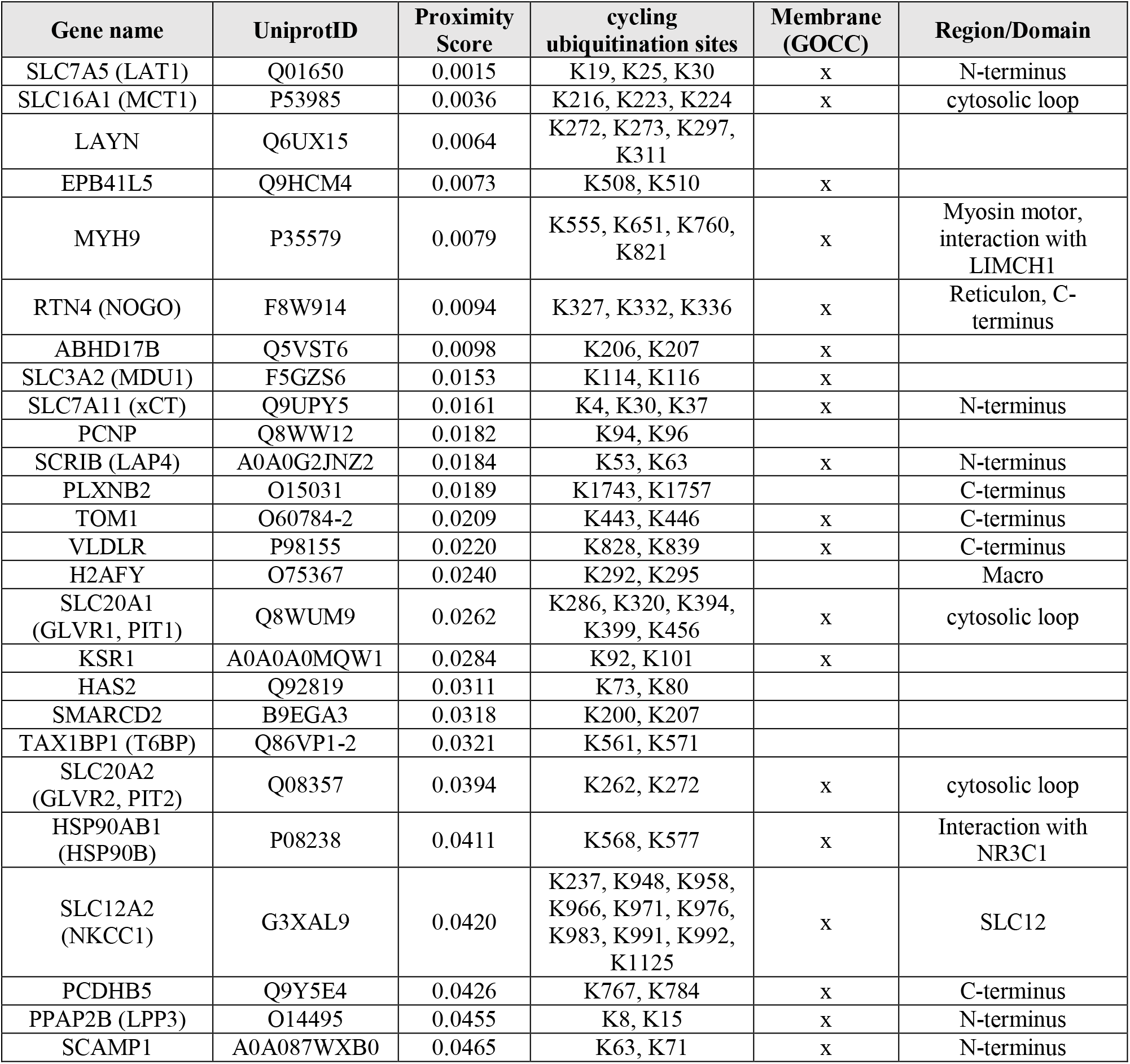
Ubiquitination clusters with potential regulatory circadian functions. Proteins with multiple cycling ubiquitination sites (q-value < 0.1) in close proximity to each other (p-value < 0.05). Membrane protein annotation by Gene Ontology Cellular Compartment (GOCC) term “membrane” and region/domain classifications are derived from UniProt and manual annotation.

Using our optimized DIA-based workflow, we identified a remarkable 33,350 ± 1,481 distinct diGly sites in single measurements of MG132 treated HEK293 samples. This implies that about half of the sites in the deep, cell line specific spectral library were matched into the single runs. Interestingly, even without using any library, a search of six single runs identified 26,800 ± 145 diGly sites (direct DIA, **Methods**). Finally, employing a hybrid spectral library - generated by merging the DDA library with a direct-DIA search - resulted in 34,733 ± 1,670 diGly sites in the same samples (**Fig. 1d**, **Supplementary Table 2).** Compared to recent reports in the literature^24^, these numbers double diGly peptide identifications in a single run format.

### DIA improves diGly proteome quantification accuracy

To evaluate the reproducibility of the entire DIA-based diGly workflow, we used MG132 treated HEK293 cells and performed three independent diGly peptide enrichments followed by DIA analysis in duplicates. This identified around 36,000 distinct diGly peptides in all replicates, 45 and 77% of which had coefficients of variations CVs below 20 and 50%, respectively (**Fig. 2a-b**, **Supplementary Table 3**). In contrast, a DDA method identified substantially fewer distinct diGly peptides and a smaller percentage with good CVs (16,000 sites; 15% with CVs < 20%; **Fig. 2a-b**). Overall, the six DIA experiments yielded almost 48,000 distinct diGly peptides, while the corresponding DDA experiments resulted in 19,000 sites.

**Fig. 2.**
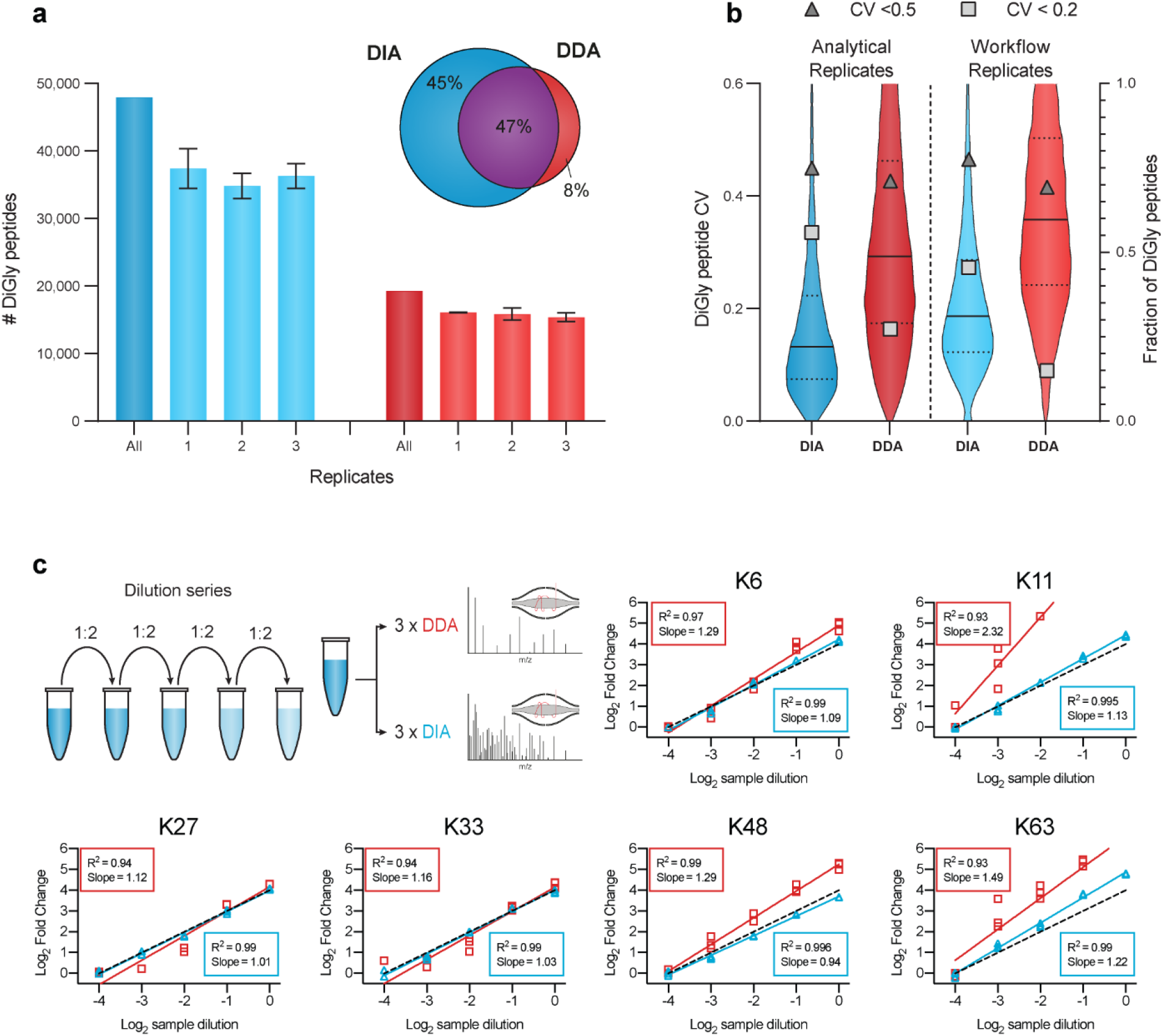
Accurate and reproducible diGly proteomics for DIA quantification. **a** Number of identified diGly peptides (± SD) for DIA (blue, HEK293 hybrid library) and DDA (red) strategies (three workflow replicates with each replicate analytical duplicates). Venn diagram depicts the proportion of shared and exclusively identified diGly sites between DIA and DDA approaches. **b** CV value distribution for DIA and DDA approaches. Solid and dotted lines denote median and 1^st^ or 3rd quantile, respectively (left axis). Fractions of CV values (right axis) below 50% and 20% are shown with filled triangles and filed squares, respectively. **c** Dilution series of diGly enriched sample. Plots show simple linear regression fits of individual ubiquitin-chain linkage type peptides measured in triplicates using DIA (blue) or DDA (red). Dotted black lines indicate the expected 1:1 slope.

To further investigate the quantitative precision and accuracy of our method, we turned to ubiquitin-chain linkage derived diGly peptides. These are the most abundant diGly peptides, all ranking in the top 20 by abundance and spanning three orders of magnitude in MS signal (**Supplementary Fig. 2g**). Diverse chain linkages confer various functions to proteins; hence, accurate quantification is important to decode the cellular roles of different ubiquitin linkage types. We performed a dilution series of a diGly sample and analyzed each dilution sample using both DIA and DDA methods in triplicates. Linear regression resulted in excellent correlations with R^2^ values higher than 0.99 for all six chain peptides assessed, much higher than the corresponding values for DDA (R^2^ 0.93-0.99; **Fig. 2c**, **Supplementary Table 3**). Importantly for quantification purposes, the experimentally observed slope for DIA was much closer to 1 than for DDA.

Together, these analytical results establish that the DIA-based workflow substantially increased the number of diGly peptides identified while markedly improving the precision and accuracy of quantification compared to a DDA-based workflow.

### In-depth ubiquitinome analysis of the TNF signaling pathway

The pro-inflammatory properties of TNF are heavily regulated by dynamic ubiquitination of its receptor-signaling complex (RSC)^32,33^ and global ubiquitnome changes upon TNF stimulation were described previously in a proteomics study^34^. Encouraged by the technical capabilities of our DIA-based diGly workflow, we here aimed to test our DIA-based diGly workflow on this well studied system, to demonstrate benefits of DIA over DDA based on accurate ubiquitination site quantification and, if possible, to extend the current knowledge of the TNF regulated ubiquitnome (**Fig. 3a**). Applying both DIA- and DDA-based diGly workflows together quantified over 10,000 diGly sites in TNF-stimulated U2OS cells (**Fig. 3b**, **Supplementary Table 4**). Both methods quantified a comparable number of ubiquitination sites (10,300 in DIA and 9,500 in DDA experiment, **Fig. 3b**). However, the DIA experiment resulted in 248 significantly upregulated ubiquitination sites (5% FDR, median fold change 2.5), of which 37 mapped to 23 proteins known to be involved in TNFα/NFκB signaling (**Fig. 3c**). In stark contrast, the DDA approach identified only 38 significant ubiquitination sites (5% FDR and median fold change 4.1), of which 15 mapped to 7 TNFα/NFκB signaling proteins. In line with these numbers, gene ontology (GO) enrichment analysis had lower FDR values and larger group sizes for terms related to the TNFα/NFκB pathway in the DIA experiment compared to DDA (**Fig. 3d**).

**Fig. 3.**
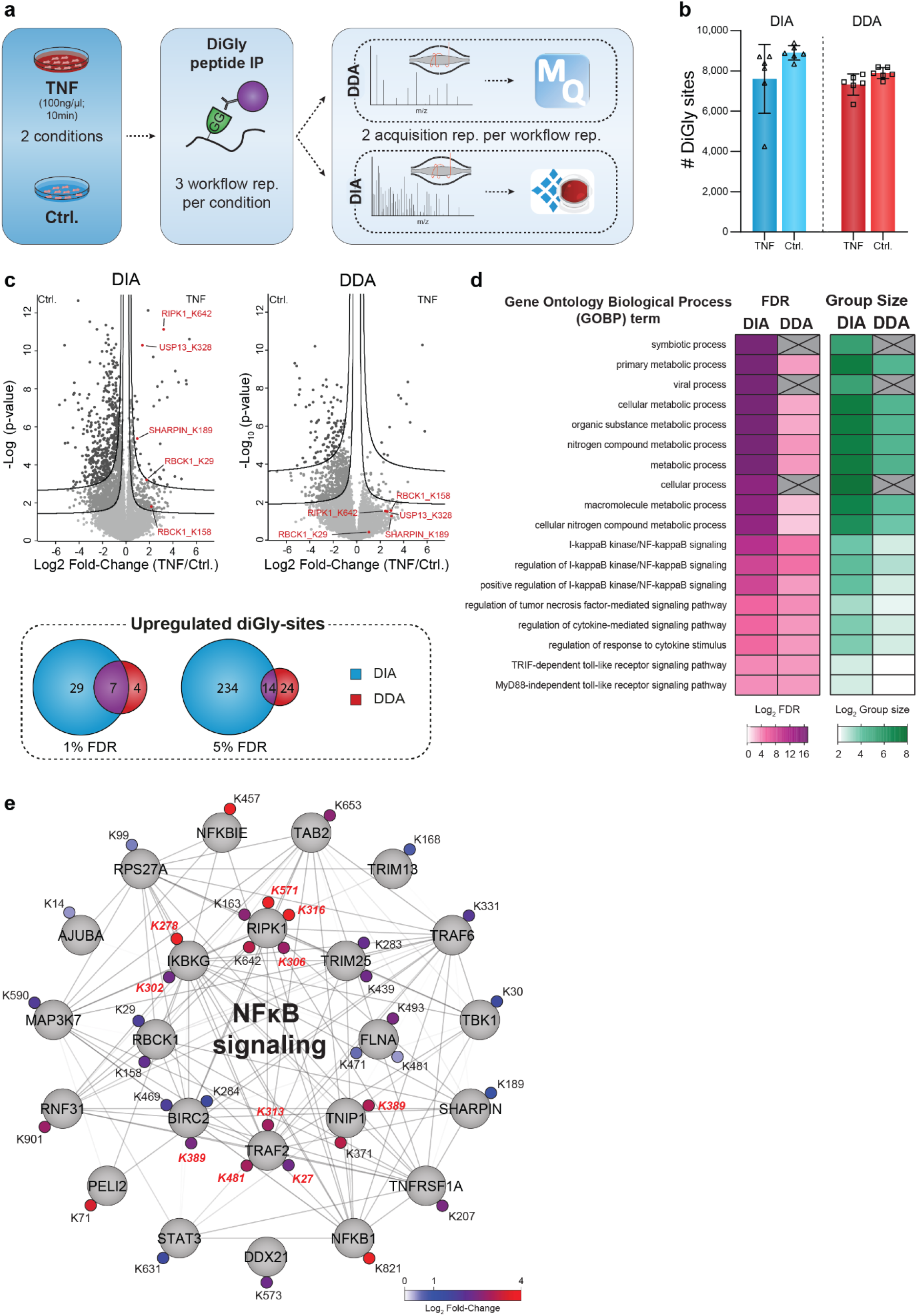
DIA enables a detailed view of the TNF regulated ubiquitnome. **a** Workflow for ubiquitnome analysis in TNF signaling. **b** Identified diGly sites (±SD) for TNF treated (10ng/μl for 10 min) and control U2OS cells in DIA (blue) and DDA (red) experiments. **c** Volcano plot of significantly regulated diGly sites at 5% FDR (lower line) and 1% (upper line) for DIA and DDA and overlaps of significantly upregulated diGly sites for 1% and 5% FDR cutoffs (t-test, s0 = 0.1). **d** Overrepresentation analysis of gene ontology biological process (GOBP) terms filtered for 5% corrected FDR (Fisher’s Exact test). **e** Cytoscape network of proteins with significantly upregulated diGly sites in DIA that are associated with NFκB signaling (GO:0043122; GO:0051092; 5% FDR). Upregulated diGly sites also captured by DDA are marked in red (5% FDR).

Several members of the TNF signaling pathway have been implicated in viral infection and TNF-receptor blockage increases susceptibility to viral infection^35,36^. The ‘viral processes’ term was significantly enriched in our DIA analysis, in line with the literature on involvement of TNF-mediated ubiquitination regulation during viral infection. Underscoring the depth of the DIA analysis, the same term failed to reach significance in the DDA analysis (**Fig. 3d**, **Supplementary Fig. 3**). In agreement with previous studies, both DIA and DDA analyses revealed increased ubiquitination of prominent members of the TNF-RSC, including TRAF2, RIPK1 and BIRC2^37,38^ (**Fig. 3e**). DIA allowed the detection of further ubiquitination events associated with the TNFα/NFκB signaling (**Fig. 3c).** For instance, the death domain (DD) of RIPK1 mediates interaction with FADD and TRADD^39^ and we found K642 in this domain to be ubiquitinated upon TNFα stimulation. Furthermore, DIA but not DDA established regulated ubiquitination of all members - HOIP/RNF31, HOIL-1/RBCK1 and Sharpin- of the LUBAC complex, a critical E3 ligase complex in TNF signaling^40,41^ in agreement with a previous study that showed LUBAC auto-ubiquitination during inflammation^42^ (**Fig. 3e**). p105/NFKB1, is a precursor for p50 and inhibitor of NFκB signaling ^43^ and we observed a striking 16-fold upregulation of K821 in its DD. Proteasome mediated limited proteolysis of p105 during NFκB signaling yields the active p50 subunit^44–47^ and the strong regulation of the K821 site suggests its involvement in this process.

DIA-based diGly analysis also uncovered TNT-regulated ubiquitination of numerous proteins known to be involved in other immune pathways. For instance, Peli2, an E3 ligase important for TLR and IL-1 signaling pathways^48^ and its interaction partner TRAF6 were ubiquitinated upon TNF stimulation. We also found that STAT2, which mediates signaling by type I interferons^49^, and USP13, which is involved in the antiviral response by deubiquitinating STING^50^, were ubiquitinated at K161 and K3218, respectively. Our results thus suggest further molecular mechanisms for the crosstalk or cross priming function by TNF to other immune pathways during viral and bacterial infections. In summary, our DIA-based ubiquitin workflow provides an in-depth view on the dynamic ubiquitination of core and peripheral members of TNF-stimulation. Apart from validating the advantages of DIA over DDA, our results provide novel regulatory ubiquitination sites, conveying a more complete picture of the various aspects of TNFα signaling.

### Circadian rhythm is globally regulated by ubiquitination

In mammals, circadian clocks are driven by interlocked transcription-translation feedback-loops. At the cellular and tissue level, they regulate oscillations of gene expression, protein abundance and post-translation modifications^51–53^. Ubiquitination plays a pivotal role in the core clock machinery (reviewed in^54^), exemplified by the ubiquitin-dependent spatiotemporal regulation of CRY proteins, the major negative clock regulators ^55^. Focused studies have provided insights into several ubiquitin-dependent events modulating core clock proteins and their effects^56–58^. Given the unexpected degree of phosphorylation-mediated signaling temporally regulated *in vivo*^51^, we wondered if ubiquitination shows similar oscillations. With the high accuracy and reproducibility of our DIA-based diGly workflow, we reasoned that it would now be possible to obtain high coverage ubiquitinome quantification across a large time series sample set to answer this question.

To this end, we measured the proteome and ubiquitinome of synchronized U2OS cells - a well-established model to study the cell autonomous circadian clock - collected every 4 hours in biological quadruplicates across 32 hours (**Fig. 4a**). Synchronization was validated by assessing the expression prolife of core clock transcripts (*Bmal1* and *Per1*) and further confirmed by PER1 and CLOCK oscillations in our proteome data (**Supplementary Fig. 5a-b**). After filtering for ubiquitinated peptides present in at least half the samples, we obtained 10,886 ubiquitination sites mapping to 3,238 proteins (**Fig. 4b**, **Supplementary Table 5**). Measurements were highly reproducible with median Pearson coefficients greater than 0.95 for biological replicates (Supplementary Fig. 5b-c). A total of 7,590 proteins were quantified in the proteome, of which at most 143 oscillated (q-value < 0.33). This small percentage of circadian regulation at the proteome level is in line with our previous proteomics results in tissues^52^ and with transcriptomics results in this cellular system^59^. Next, we normalized the intensities of the diGly peptides encompassing each ubiquitination site to their corresponding protein abundance. The resulting quantitative values represent the occupancy of the ubiquitin sites irrespective of changes in protein abundance (**Methods**, **Supplementary Fig. 5d**).

**Fig. 4.**
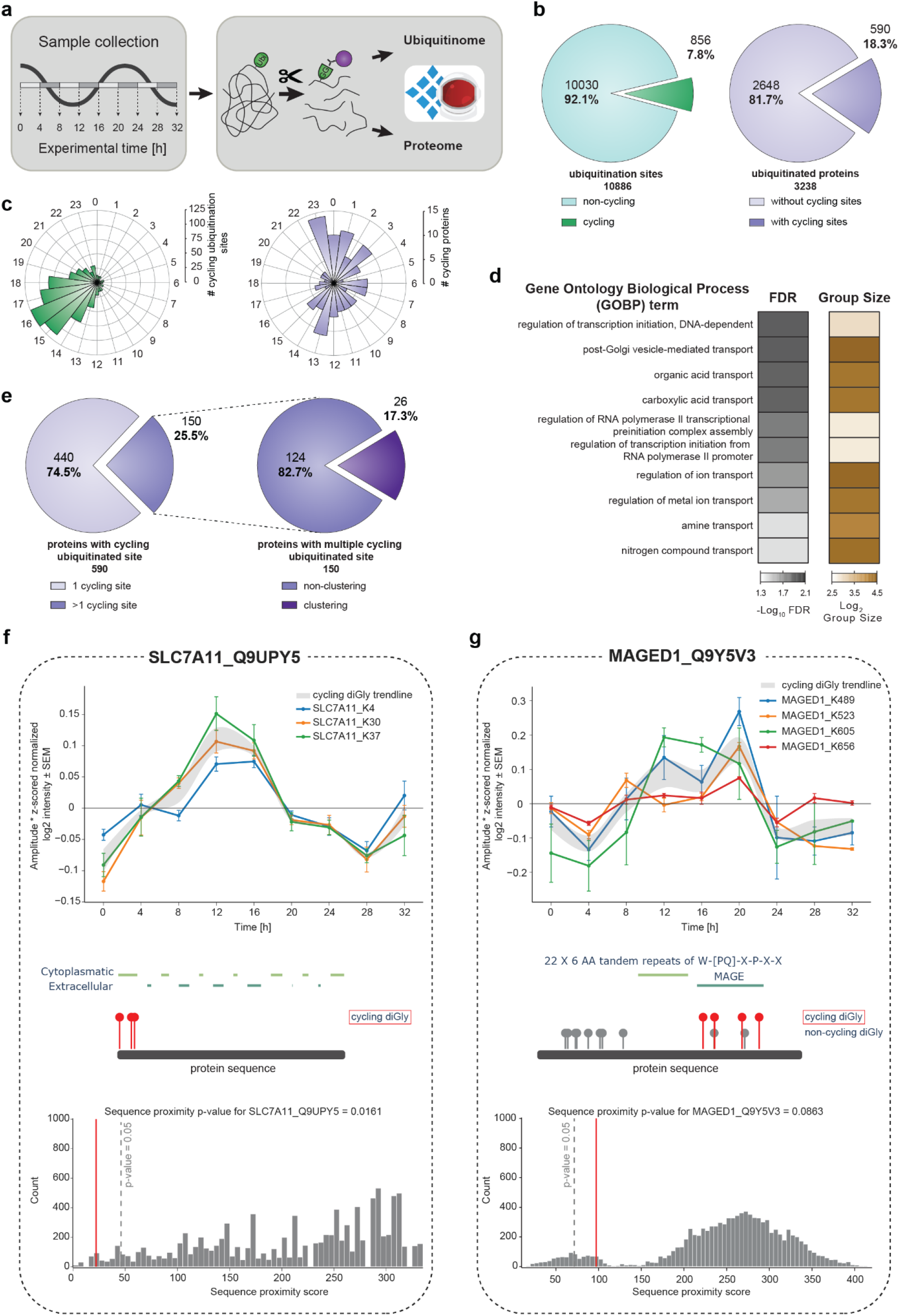
Quantification of the rhythmic ubiquitinome. **a** Experimental workflow for rhythmic ubiquitinome analysis. **b** Proportion of oscillating ubiquitination sites (q-value < 0.1) quantified in > 50% of all samples (left panel) and proteins with cycling ubiquitin sites (q-value < 0.1) (right panel) **c** Rose plots indicate phase peaks for cycling ubiquitination sites (left panel) and proteins (right panel). **d** Overrepresentation analysis of gene ontology biological processes (GOBP) filtered for top 10 significant terms. Significance is determined by 5% corrected FDR (Fisher’s Exact test). **e** Proportions of proteins with a single and multiple cycling ubiquitination sites (left panel) and those displaying cycling diGly site clusters (right panel). Examples of proximity analysis of cycling ubiquitin clusters (http://cyclingubi.biochem.mpg.de). Cycling sites (q-value < 0.1) (top) and their location in the protein sequence along with the domain annotation (middle) and proximity score (average distance, p-value < 0.1) (bottom) for SLC7A11 **f** and MAGED1 **g**.

Periodicity analysis showed that 8% of the ubiquitination sites on 18% of the proteins oscillated in a circadian manner (856 sites; 590 proteins, **Methods**, q-value < 0.1, **Fig. 4c** and **Supplementary Fig. 5e**). A large proportion of rhythmic sites peaked with phases clustered around 16-20 hours after synchronization (**Fig. 4c**, **Supplementary Fig. 5e**). Remarkably, 59% of these were annotated to be in membrane proteins, many more than expected by chance (p < 10^−172^; **Supplementary Table 5**). Overrepresentation analysis revealed that these proteins are predominantly involved in transport of a small molecules, such as ions, amines and organic acids (**Fig. 4d**). These findings point to a potential metabolic function of circadian membrane protein ubiquitination.

A full quarter of rhythmic ubiquitinated proteins harbored more than one oscillating site (150 sites; **Fig. 4e**). To investigate the spatial arrangement of them, we developed a bioinformatic proximity analysis tool (available as part of our website for browsing and analyzing the cellular ubiquitinome http://cyclingubi.biochem.mpg.de). In 17% of proteins rhythmic ubiquitination sites were closer together than expected by chance (p < 0.05) and 73% were annotated as membrane proteins. Interestingly these adjacent sites were mostly located in regions with potential regulatory function, such as N- and C-termini, cytosolic loops and interaction domains (**Table 1**). For instance, K4, K30 and K37 of the sodium independent cystine-glutamate transporter (SLC7A11, 501 aa) are rhythmically ubiquitinated with similar phases (13.8; 13.3; 13.1 h, respectively, **Fig. 4f**). Likewise, the potassium chloride symporter NKCC1 (SLC12A2) has a cluster of eight rhythmically ubiquitinated sites in its C-terminal domain with similar phases (K948, K958, K966, K971, K976, K983, K991, K992; **Supplementary Fig. 5f**). This widely expressed solute carrier plays a key role in the regulation of ionic balance and cell volume^60^. We also discovered novel oscillating ubiquitin modifications in the MAGE domain of MAGED1, a protein that directly interacts with the core clock protein RORα, to regulate Bmal1, Rev-erbα and E4bp4 gene expression (**Fig. 4g**). Interestingly, despite these rhythmic outputs neither the *Maged1* transcript, protein expression nor its binding to RORα oscillates^61^. Our results now suggest that MAGED1 activity could instead be rhythmically controlled post-translationally through the multiple ubiquitinations in its MAGE domain.

Together, this first in-depth view of the circadian ubiquitinome, made possible by our DIA-based diGly workflow, reveals this PTM to be a major regulatory mechanism driving rhythmic processes, which include essential cellular processes such as ion transport and osmotic balance.

## DISCUSSION

We here developed a sensitive and robust DIA-based workflow, capable of identifying 35,000 diGly peptides in single run measurements. Both the depth of coverage and the quantitative accuracy are unprecedented and are doubled compared to otherwise identical DDA experiments. Importantly the workflow requires no extra labeling step or offline fractionation, making it streamlined and easy to implement. Furthermore, it could be used for quantification of other PTMs relying on antibody-based enrichment such as lysine acetylation and tyrosine phosphorylation. A current limitation of the DIA method is that, like for any DIA-based analysis, including phosphoproteome analysis ^25,30^, the best coverage and quantification is obtained with custom-made, project-specific spectral libraries. Construction of such spectral-libraries requires some effort, specialized equipment for fractionation and may not always be possible for samples with low amounts such as primary cells. Library-free approaches would greatly aid to simplify DIA workflows in the future and there are considerable efforts currently being invested into producing prediction tools for MS/MS spectra and retention time^62^.

Compared to single run TMT-based workflows, DIA suffers from lower throughput. However, the latest advances in nanoflow liquid chromatography now increasingly allow rapid, robust and deep DIA-based proteome and phosphoproteome profiling, which is likely applicable to DIA-based ubiquitinome analysis as well. Furthermore, the LC MS/MS analysis of our workflow requires only a few hundred μg and it already enables the analysis of systems such as human primary cell culture models where protein material is limited. However, further sensitivity advances are limited by the initial antibody-based enrichment, which currently requires 0.5-1 mg of sample. If this step could be scaled down and the subsequent peptide purification eliminated altogether, sample size requirements could become much smaller yet. A workflow without a peptide-clean-up step would also aid to further improve throughput and reproducibility, making the entire workflow more streamlined.

By converting from a DDA to a DIA workflow we demonstrate a dramatic increase in the number of ubiquitination sites that can consistently and significantly be quantified. Given the inherent sensitivity of our single run approach allowing system-wide investigations of ubiquitination dynamics of biological processes, we applied it to TNF signaling. This provided an in depth view on the ubiquitination dynamics of TNF signaling, covering core and peripheral signaling members, which a parallel DDA analysis failed to provide. Apart from validating advantages of DIA over DDA, our results show that, like phosphorylation, ubiquitination signaling events are rapidly induced after TNF stimulation. Unexpectedly, we still pinpointed novel TNF-regulated sites on proteins that were not previously described in this well-studied pathway. The rich resource provided here could be further explored to investigate the functions of these ubiquitination events in TNFα signaling in health and disease.

System-wide circadian proteomics studies have so far been limited to the dynamic regulation of protein and phosphorylation levels – largely for technological reasons. Our in-depth quantitative diGly analysis of the cell autonomous circadian clock now extends those studies by providing the first cell intrinsic circadian map of ubiquitination dynamics. Quantifying more than 10,000 unique ubiquitination sites in synchronized U2OS cells, a standard cellular model of the circadian rhythm, revealed that 8% of them - located on 18% of the quantified ubiquitinated proteins - oscillated in abundance. Many of the cycling sites match into the DIA library of untreated, rather than the library of proteasome inhibited cells suggesting they could have regulatory, non-degradative functions.

Our data reveal wide-spread ubiquitination of membrane proteins, transporters and receptors, proteins that regulate major cellular processes such as cell volume, ion balance and osmotic homeostasis. Intriguingly, often these cycling ubiquitination sites on membrane proteins are not randomly distributed over the protein sequence but rather cluster in certain regions such as the N- and C-terminus. Circadian rhythms in Mg2+ and K+ cellular levels and their transport have been reported in a range of eukaryotic cell types suggesting an evolutionary conservation of this mechanism. Moreover, K+ transport is a key mechanism driving electrical excitability oscillations in the mammalian master clock and Drosophila neurons ^63,64^ and in turn, plasma membrane potential feeds back to the cellular clock ^65,66^. Despite their fundamental cellular role, little is known about the regulatory mechanisms controlling rhythms of ion levels and size in cells ^67,68^. Our system-level data suggest that ubiquitination plays a major role in the rhythmic transport of ions and other compounds in the cell by temporally modulating the activity of membrane transporters. Such a mechanism would, for instance, explain the observation that red blood cells lose their daily electrophysiological rhythm after proteasome treatment^67^.

We speculate that ubiquitin-dependent temporal regulation of transporter function for various substrates (e.g. sodium/phosphate/chloride - SLC20A1/SLC20A2, monocarboxylates - SLC16A1, Sodium/Potassium - ATP1A1, various amino acids - SLC3A2, SLC7A5, SLC7A11, and organic anions - ABCC3) and other receptors (e.g. TGFBR2 or PLXNB2) may serve as temporal cellular switches to sense and respond to daily changes in nutrient availability. Interestingly, in our recent phosphoproteomics study of the synaptic compartment we observed that many of the ubiquitination-related proteins had rhythmic phosphorylation sites^69^. This suggests an interplay between post-translational modifications that together could fine tune daily cycles of membrane-mediated processes essential for proper cellular and tissue metabolism. Given the central role of transporters in chronopharmacology70–72, ubiquitin-dependent dynamic regulation of specific membrane transporters are an important functional aspect to consider for drug administration and patient health, both key goals of chronotherapy. The data of our rhythmic ubiquitinome analysis is accessible at http://cyclingubi.biochem.mpg.de, opening up new avenues for mechanistic investigations.

## ACKNOWLEDGEMENT

FB and MSR were supported by the Volkswagen Foundation (93071), MSR also received funding from the German Research Foundation DFG (Project 329628492-SFB1321, INST 86/1800-1 FUGG and project 428041612). We thank all the members of the Department of Signal Transduction and Proteomics, in particular, Igor Paron and Christian Deiml for MS technical assistance, Bianca Splettstoesser for technical help with experimental work and Johannes Müller, Alex Strasser and Lisa Schweitzer for columns. We also thank Sharah Meszaros and Steve Dewitz for cell culture assistance. We are grateful to Arno F. Alpi, Jesper Olsen, Chuna Choudhary, André Michaelis and Jakob Bader for constructive and insightful discussions. We specially thank Florian Gnad and Cell Signaling Technologies (CST) for gifting of PTMScan® Ubiquitin Remnant Motif (K-ɛ-GG) Kits.

## AUTHOR CONTRIBUTIONS

FMH, OK, MT, FB and MSR designed experiments. TNF experiments were performed by FMH and MT. Circadian experiments were conducted by FMH and FB. Computational proximity analysis was performed by IB. Data was analyzed by FMH. FMH, OK, MT, MSR, BAS and MM wrote, reviewed and edited the manuscript. All authors read and commented on the manuscript.

## METHODS

### Cell culture, treatment, harvest and lysis

HEK293 (human, DMSZ, ACC 635) and U2OS (human, American Type Culture Collection [ATCC], HTB-96) cell were cultivated in DMEM (Gibco, Invitrogen) supplemented with 10% fetal bovine serum (Gibco, Invitrogen), 100 U/ml penicillin (Gibco, Invitrogen) and 100 μg/ml streptomycin (Gibco, Invitrogen) at 37°C in a humidified incubator with a 5% CO_2_ atmosphere. For cell harvest, cells were washed twice with ice-cold PBS (Gibco, Invitrogen), centrifuged, snap frozen in liquid nitrogen and stored at −80°C until lysis. Frozen cell pellets were lysed by adding lysis buffer (1% SDC in 100 mM Tris/HCl, pH 8.5) directly onto frozen cell pellets, followed by repeated aspiration and boiling for 5 min at 95°C.

For proteasome inhibition, HEK293 or U2OS cells were treated with 10 μM MG132 (InvivoGen) at approximately 80% confluence for 4 h and successively harvested. For circadian cycle experiments, cells were synchronized, when they reached at least 90% confluence, with dexamethasone (1 μM) for 1 h. Following this, U2OS were washed once with PBS and the medium was replaced. The first time point was collected after 24 hours of synchronization continuing the collection every 4 hours across 32 hours for each of the 4 biological replicates. Collected cells, stored and lysed as described. For TNF stimulation of U2OS cells, confluent cultures were either stimulated with 100 ng/ml TNF for 10 minutes or left unstimulated. Cells were washed 3 x with ice-cold PBS, directly lysed with lysis buffer and boiled for 5 min at 95°C.

### Western blot analysis

U2OS cells were plated in 6 well plates and when confluent stimulated for 5, 10, 15, 30, 60 minutes with 100 ng/ml TNF or left untreated. After stimulation, cells were washed in PBS and lysed in 4% SDS in 100 mM Tris/HCL, pH 8. Lysates were boiled, sonicated and protein concentrations were estimated using BCA. SDS sample loading buffer (450 mM Tris-HCl, pH 8, 60% (v/v) glycerol, 12% (w/v) SDS, 0.02% (w/v) bromophenol blue, 600 mM DTT) was added to lysates before separation on 12% Novex Tris-glycine gels (Thermo Fisher Scientific, XP00120BOX). Separated proteins were transferred onto PVDF membranes (Merck Millipore, IPVH00010). Membranes were blocked in 5% BSA in PBST and antibodies diluted in 2% BSA in PBST. Antibodies used for immunoblotting were as follows: anti phospho p65 (CST, 3033P), anti p65 (CST, 4764P), anti IκBα (CST, 92424792), anti phospho p38 (CST, 9215), anti p38 (CST, 9212), anti β-actin (CST, 4970).

### RNA isolation and QPCR

RNA was isolated from 3 biological replicates of each U2OS time point according to manufacture instruction using the RNeasy Plus Mini Kit (QIAGEN, #74134). Isolated RNA was reversely transcribed by using first-strand cDNA synthesis kit (Thermo Fisher Scientific, #K1612). QPCR was performed at the C1000 Thermal Cycler (Bio-Rad) with iQ™ SYBR Green Supermix (Bio-Rad, #170-8862) with following primers:

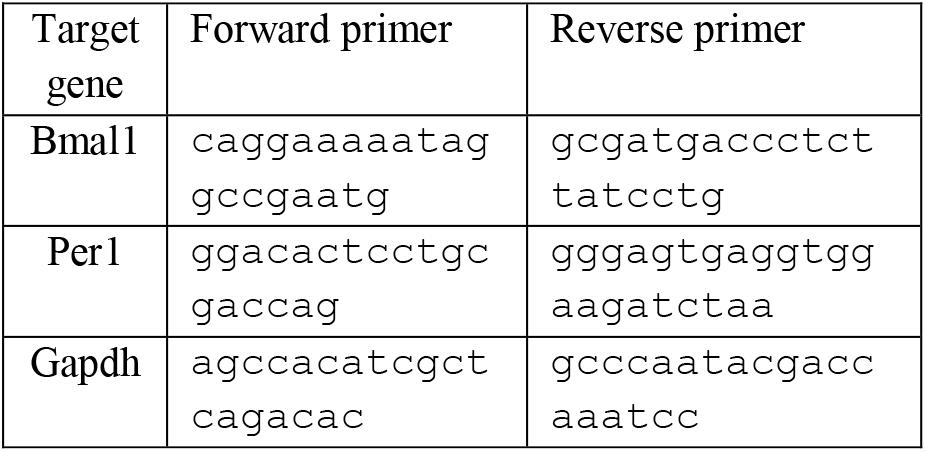

The in-build analysis tool of the CFX Manager Software (Version 3.1, Bio-Rad) was used to determine the normalized expression with the ΔΔCq method of Bmal1 and Per1 compared to Gapdh in technical triplicates for all 3 biological replicates of each time point. The technical triplicates were further averaged and adjusted so that the highest value was set to 1. Following this, the average of all biological replicates and the SEM (standard error of the mean) was calculated for all the time points.

### Protein digestion and peptide cleanup

Lysates were sonicated for 1 min (Branson Sonifier) and protein concentrations were estimated by tryptophan assay. After addition of CAA and TCEP to a final concentration of 10 mM and 40 mM, respectively, samples were incubated for 5 min at 45°C for protein reduction and alkylation. Thereafter, Samples were digested overnight at 37°C using trypsin (1:100 w/w, Sigma-Aldrich) and LysC (1/100 w/w, Wako).

For proteome analysis, sample aliquots (~15μg) were desalted in SDB-RPS StageTips (Empore). Briefly, samples were first diluted with 1% TFA in isopropanol to a final volume of 200 μl. Thereafter, samples were loaded onto StageTips and sequentially washed with 200 μl of 1% TFA in isopropanol and 200 μl 0.2% TFA/ 2% ACN. Peptides were eluted with 60 μl of 1.25% Ammonium hydroxide (NH_4_OH)/ 80% ACN and dried using a SpeedVac centrifuge (Eppendorf, Concentrator plus). Dried peptides were resuspended in buffer A* (2% ACN/ 0.1% TFA) supplemented with iRT peptides (1/30 v/v) (iRT Standard, Biognosys).

For diGly peptide enrichment, samples were four-fold diluted with 1% TFA in isopropanol and loaded onto SDB-RPS cartridges (Strata™-X-C, 30 mg/ 3 ml or Strata™-X-C, 200 mg/ 6 ml, Phenomenex Inc.). Before peptide loading, cartridges were equilibrated with 8 bed volumes (BV) 30% MeOH/1% TFA and washed with 8 BV of 0.2% TFA. Samples were loaded by gravity flow and sequentially washed twice with 8 BV 1% TFA in isopropanol and once with 8 BV 0.2% TFA/ 2% ACN. Peptides were eluted twice with 4 BV 1.25% NH_4_OH/ 80% ACN and diluted with ddH_2_O to a final ACN concentration of 35% ACN. Thereafter, samples were snap frozen in liquid nitrogen, lyophilized and store at 4°C until diGly peptide enrichment.

### DiGly peptide enrichment

Lyophilized peptides were resuspended in immunoaffinity purification buffer (IAP) (50 mM MOPS, pH 7.2, 10 mM Na_2_HPO_4_, 50 mM NaCl) and sonicated for 2.5 min (Bioruptor plus, Diagenode). Peptide concentration was estimated by tryptophan assay. DiGly remnant containing peptides were enriched using the PTMScan® Ubiquitin Remnant Motif (K-ɛ-GG) Kit (Cell Signaling Technology (CST)), which was kindly provided by CST. First, antibodies were crosslinking to beads as described by Udeshi et al.^22^. Unless otherwise stated, 1/8 of a vial of crosslinked antibody beads and 1 mg of peptide material were used for diGly peptide enrichments. For this, peptides were added to crosslinked antibody beads and the volume was adjusted to 1 ml with IAP buffer. After 1h of incubation at 4°C and gentle agitation, beads were washed twice with cold IAP and 5 times with cold ddH_2_O. For this, beads were transferred into GF-filter StageTips and for each wash step, the according wash solution was added and passed through by centrifugal force. Thereafter, GF-StageTips were stacked onto SDB-RPS StageTips and peptides were directly eluted into SDB-RPS StageTips. For this, 50 μl 0.15% TFA were added twice onto the beads and passed through by centrifugation for 5 min at 100 g. Thereafter, 100 μl 0.2% TFA were added on top of peptide eluates, followed by sample loading onto the stationary material of SDB-RPS StageTips. Peptides were washed, eluted and dried as described for proteomes samples, with the difference, that 0.2% TFA was used for the first wash step. Dried peptides were resuspended in 9 μl buffer A*, supplemented with iRT peptides (1/30 v/v) for LC/MS-MS analysis.

### Basic reversed phase fractionation

Basic reversed phase (bRP) fractionation for diGly peptide and proteome spectral libraries were performed on an UFLC system (Shimadzu) and EASY-nLC 1000 (Thermo Fisher Scientific, Germany), respectively.

For diGly peptide separation, lyophilized samples were resuspended in Buffer A (5 mM NH_4_HCO_2_/ 2% ACN) and 5 mg peptide material (5 mg/ml) was loaded onto a reversed phase column (ZORBAX 300Extend-C18, Agilent). Peptides were separate at a flow rate of 2 ml/min and a constant column temperature of 40 °C using a binary buffer system, consisting of buffer A and buffer B (5 mM NH_4_HCO_2_/ 90% ACN). An elution gradient staring at 0% B stepwise increased to 28 in 53 min and to 78 in 6 min was deployed. Eluting peptides were automatically collected into a 96-deepwell plate while well positions were switched in 40 s intervals.

For peptide fractionation on the EASY-nLC 1000 system, approximately 55 μg peptide material were loaded onto a 30 cm in-house packed, reversed phase columns (250 μm inner diameter, ReproSil-Pur C18-AQ 1.9 μm resin [Dr. Maisch GmbH]). Peptides were separate at a flow rate of 2 μl/min using a binary buffer system of buffer A (PreOmics) and buffer B (PreOmics). An elution gradient staring at 3% B stepwise increased to 30% in 45 min, 60% in 17 min and 95% in 5 min was used. Eluting peptides were concatenated into 24 fractions by switching the rotor valve of an automated concatenation system (Spider fractionator, PreOmics) ^73^ in 90 s intervals.

### Library sample preparation

For individual deep diGly libraries 2×5 mg peptide were fractionated by bRP fractionation. For K48-peptide containing fraction identification, 100μl aliquots of fractions 46 to 54 were dried in a SpeedVac, resuspended in A* and measured on an LTQ Orbitrap XL mass spectrometer. K48-peptide containing fractions of both plates, were pooled in sample pool “K48” (**Supplementary Fig. 1a**). Remaining fractions of both pates were concatenated into P1-P8 (**Supplementary Fig. 1a**), snap frozen and lyophilized. Lyophilized peptides were resuspended in 1 ml IAP buffer and diGly peptides were enriched as described above. In case of HEK293 library generation, an optional second supernatant IP was conducted. For this, 500μl of previous diGly peptide enrichment supernatants were pooled as indicated (**Supplementary Fig. 1a**) and used for sequential diGly peptide enrichment.

For the proteome library, aliquots of U2OS samples for proteome cycling analysis were used. Approximately 3 μg peptide material of individual time points of two biological replicates, after SDB-RPS cleanup, were pooled and fractionate via bRP fractionation as described above. Fractionated samples were dried using a SpeedVac and resuspended in A* supplemented with iRT peptides (1/30 v/v) for LC-MS/MS measurement and spectral library generation.

### Nano-flow LC-MS/MS proteome measurements

Peptides were loaded onto a 50 cm, in-house packed, reversed phase columns (75 μm inner diameter, ReproSil-Pur C18-AQ 1.9 μm resin [Dr. Maisch GmbH]). The column temperature was controlled at 60°C using a homemade column oven and binary buffer system, consisting of buffer A (0.1% formic acid (FA)) and buffer B (0.1% FA in 80% ACN), was utilized for low pH peptide separation. An EASY-nLC 1200 system (Thermo Fisher Scientific), directly coupled online with the mass spectrometer (Q Exactive HF-X, Thermo Fisher Scientific) via a nano-electrospray source, was employed for nano-flow liquid chromatography, at a flow rate of 300 nl/min. For individual measurements, 500 ng of peptide material was loaded and eluted with a gradient starting at 5% buffer B and stepwise increased to 30% in 95 min, 60% in 5 min and 95% in 5 min.

The same general setup was used, for K48-peptide containing fraction identification, while the column and mass spectrometer were changed to a 20 cm column and an LTQ Orbitrap XL, respectively.

For DDA experiments the Thermo Xcalibur (4.0.27.19) and LTQ Tune plus (2.5.5 SP2) software were used for Q Exactive HF-X and LTQ Orbitrap XL instruments, respectively. The Q Exactive HF-X was operated in Top12 mode with a full scan range of 300-1650 *m/z* at a resolution of 60,000. The automatic gain control (AGC) was set to 3e6 at a maximum injection time of 20 s. Precursor ion selection width was kept at 1.4 *m/z* and fragmentation was achieved by higher-energy collisional dissociation (HCD) (NCE 27%). Fragment ion scans were recorded at a resolution of 15,000, an AGC of 1e5 and a maximum fill time of 60 ms. Dynamic exclusion was enabled and set to 20 s. The LTQ Orbitrap XL was operated in Top10 mode with a full scan range of 300-1700 *m/z* at a resolution of 60,000. Precursor ion selection width was kept at 2.0 *m/z* and fragmentation was achieved by collision-induced dissociation (CID) (NCE 35%).

For DIA analysis, the MaxQuant Live software suite was utilized for data acquisition^74^. The full scan range was set to 300-1650 *m/z* at a resolution of 120,000. The AGC was set to 3e6 at a maximum injection time of 60 ms. HCD (NCD 27%) was used for precursor fragmentation and fragment ions were analyzed in 33 DIA windows at a resolution of 30,000, while the AGC was kept at 3e6.

### Nano-flow LC-MS/MS diGly measurements

DiGly peptide enriched samples were measured on a Q Exactive HF-X using the same instrumental setup as for proteome analysis. For diGly single run measurements one quarter (2 μl) and for diGly library preparation one-half (4 μl) of enriched samples were loaded for LC-MS/MS analysis, unless stated otherwise. Loaded peptides were eluted using a gradient starting at 3% buffer B and stepwise increased to 7% in 6 min, 20% in 49 min, 36% in 39 min, 45% in 10 min and 95% in 4 min.

For DDA analysis, the MS was operated in Top12 mode with a full scan range of 300-1350 *m/z* at a resolution of 60,000. AGC was set to 3e6 at a maximum injection time of 20 s. Precursor ion selection width was kept at 1.4 *m/z* and fragmentation was achieved by HCD (NCE 28%). Fragment ion scans were recorded at a resolution of 30,000, an AGC of 1e5 and a maximum fill time of 110 ms. Dynamic exclusion was enabled and set to 30 s.

For DIA analysis, the MaxQuant Live software suite was employed for data acquisition^74^. The full scan range was set to 300-1650 *m/z* at a resolution of 120,000. The AGC was set to 3e6 at a maximum injection time of 60 ms. HCD (NCD 28%) was used for precursor fragmentation and resulting fragment ions were analyzed in 46 DIA windows at a resolution of 30,000 (unless otherwise stated) and an AGC of 3e6. DIA window distribution parameters PdfMu and PdfSigma were set to 6.161865 and 0.348444, respectively, unless stated otherwise.

### Raw data analysis

DDA raw data used for K48-peptide fraction identification and DIA and DDA comparisons were analyzed with MaxQuant (1.6.2.10) using default settings and enabled match between runs (MBR) functionality. Carbamidomethyl (C) was defined as fixed modification and Oxidation (M), Acetyl (Protein N-term) and DiGly (K) were set as variable modifications.

DDA raw data, used for spectral library construction, were processed with Spectronauts build in search engine pulsar (13.12.200217.43655)^28^. Default settings were used for proteome spectral libraries. For diGly spectral libraries the “Best N Fragments per peptides” maximum value was adjusted to 25. For hybrid library construction DIA raw files were processed together with DDA library raw files using the same search settings.

DIA raw files were processed using Spectronaut (13.12.200217.43655)^28^. Proteome analysis was performed with default settings. For diGly analysis, diGly (K) was defined as an additional variable modification and PTM localization was enabled and set to 0. For dilution experiments, “XIC RT extraction window” was set to “static” with a window width of 10 min. Direct DIA searches used the same settings as described above.

### Bioinformatics analysis

Data analysis was primarily performed in the Perseus software suite (1.6.7.0). For diGly site analysis, Spectronaut normal report output tables were aggregated to diGly sites using the peptide collapse plug-in tool for Perseus^30^. DiGly sites were aggregated using the linear model based approach and filtered for a localization probability > 0.5. Student t-test statistics (FDR cutoff 1% or 5%; s0 = 0.1) for TNF stimulation experiments were performed in Perseus. Fisher’s Exact GOBP Term enrichment of upregulated diGly sites and cycling diGly sites was performed on the pantherdb website (http://pantherdb.org/) and in perseus, respectively, with Benjamini Hochberg FDR correction enabled and set to a 5% cutoff. Network representation of upregulated diGly sites was performed with the STRING app (1.5.1) in Cytoscape (3.7.2).

For the cycling analysis of diGly sites, data was first filtered for diGly sites identified in at least 50% across all measurements. Proteins and diGly sites raw intensities were log_2_ transformed and normalized by median subtraction. For diGly site protein normalization the median values of biological quadruplicates were subtracted from normalized diGly sites. Missing values of protein data for subtraction were imputed based on a Gaussian normal distribution with a width of 0.3 and a downshift of 1.8. Cycling analysis of normalized protein and diGly site data was performed as previously described, but in this case with a period time of 24.8 h^51,52^. A q-value cutoff of < 0.1 and < 0.33 was used to define cycling DiGly sites and proteins, respectively.

### Website tool

For profile plots individual z-scores for each protein abundance normalized diGly site and the median z-score and standard error of means (SEM) were subsequently determined for each time point. The resulting median z-scores and SEM values were multiplied with the cycling amplitude of each diGly site (Perseus periodicity analysis output). For sequence visualization and protein domain annotation each diGly site location was mapped to the first UniProt ID of its assigned protein group and was visualized based on its respective protein sequence stored in the fasta file that was used for MS/MS data analysis (human fasta, downloaded 2015). The protein sequences for visualization were obtained using the ‘fasta’ functions from pyteomics^75,76^. Information about protein domains was obtained from UniProt (https://www.uniprot.org/, accessed 25.05.2020), including following categories: ‘Topological domain’, ‘Motif’, ‘Motif’, ‘Motif’, ‘Zink finger’ and ‘Domain [FT]’.

To evaluate whether multiple observed cycling diGly sites are located in a specific region on the protein, we performed a proximity analysis. Three different metrics were evaluated: (1) the average distance (In amino acids) between all observed cycling diGly sites, (2) the minimum distance between any two observed cycling diGly sites, and (3) the maximum distance between any two observed cycling diGly sites. The observed distance metrics were compared to the distances expected from a random distribution of the diGly sites of a protein across all of its lysines. 10,000 random distributions were considered and an empirical p-value was estimated based on the fraction of random samples with a smaller or equally small distance metric as the observed cycling diGly sites. For the main analysis, diGly sites with a q-value ≤ 0.1 were considered as cycling diGly sites.

Data preprocessing and visualization for the dashboard was performed using the python programming language. Following libraries were utilized for data processing: numpy, pandas, re, random and pyteomics ^75,76^. Several libraries from the HoloViz family of tools were used for data visualization and creation of the dashboard, including panel and holoviews, but also bokeh, plotly and matplotlib.

## Supplementary Figures

**Supplementary Figure 1.**
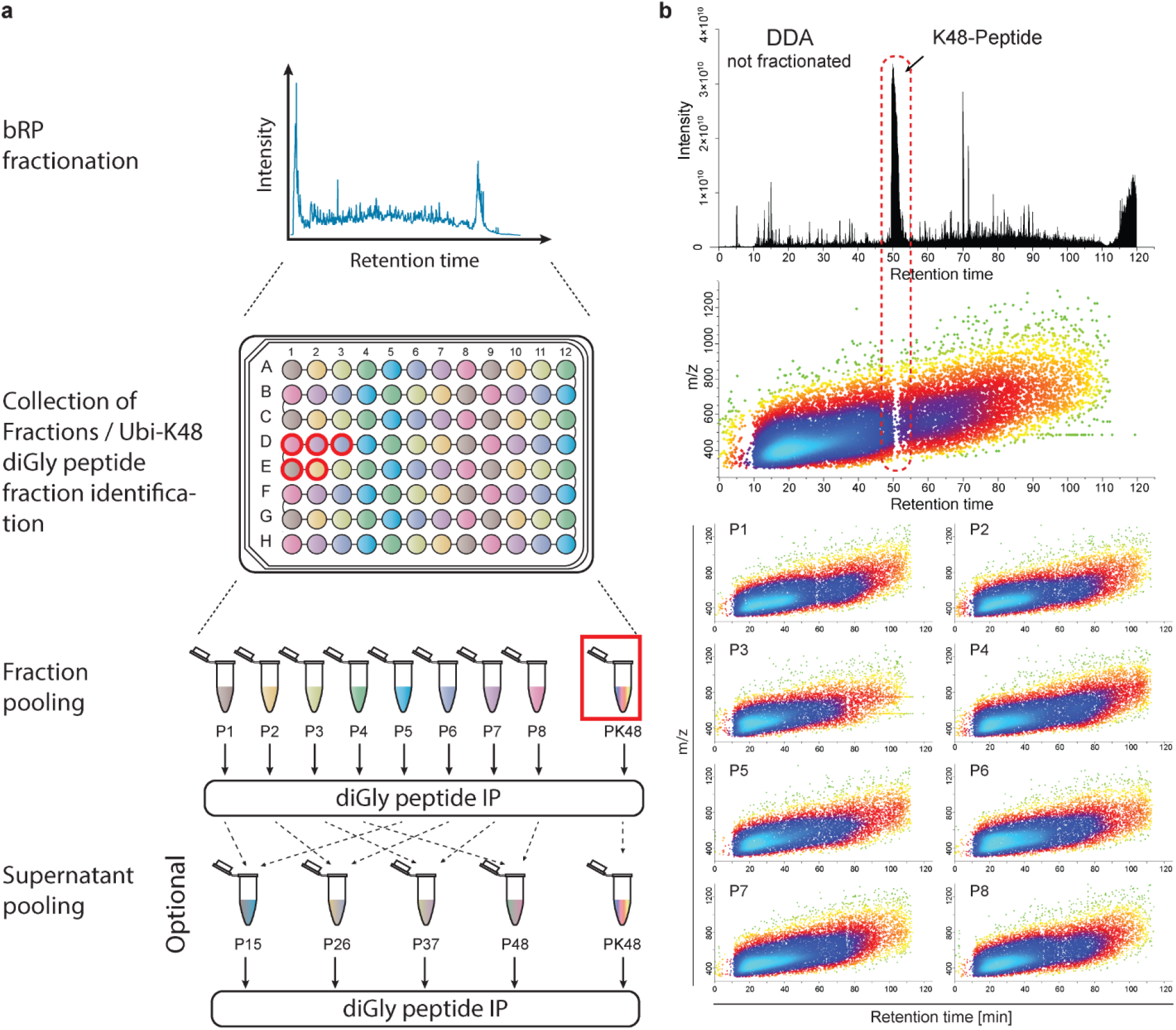
DiGly sample fractionation and concatenation scheme. **a** Workflow for bRP sample fractionation, K48-peptide fraction exclusion and sample concatenation. **b** Characteristic chromatogram and density plot for diGly analysis via DDA (upper panels) and density plots for individual sample pools (P1-P8).

**Supplementary Figure 2.**
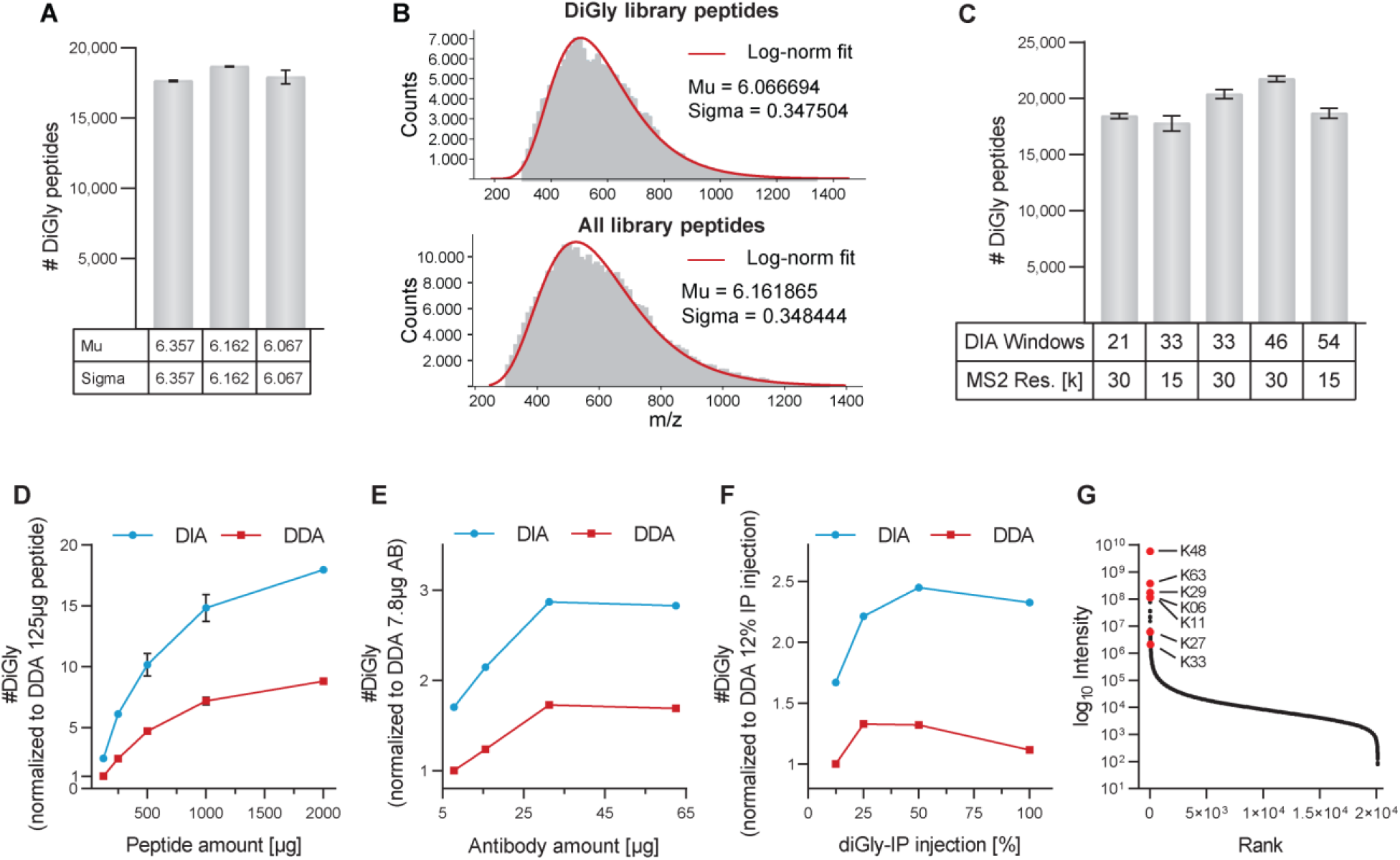
DIA method optimization. **a** Number of identified diGly peptides (±SD) for MaxQuant Live input value variation of *m/z* peptide distribution log-Norm fit values: Mu and Sigma. MaxQuant Live default values (left), log-Norm fit values of all identified peptides (middle) and only diGly peptides (right) of HEK293 library peptides. **b** Lognormal fit curves for diGly (upper panel) and all identified peptides (lower panel) in HEK293 library. **c** DiGly peptide identification (±SD) for different DIA window number and MS2 resolution setting. **d** Relative diGly site identification (±SD) of different peptide starting amounts using 31.25 μg antibody for enrichment. **e** Same as **d** with constant peptide input (1mg) and varying antibody amounts. **f** Same as **d** with constant peptide (1mg) and antibody amount (31.25 μg) with varying sample injection amounts **g** Dynamic range of diGly peptides. Red dots highlight ubiquitin chain linkage derived diGly peptides. Validation of TNFα signaling induced upon TNF stimulation. U2OS cells were stimulated with 100 ng/μl TNF for the indicated time.

**Supplementary Figure 3.**
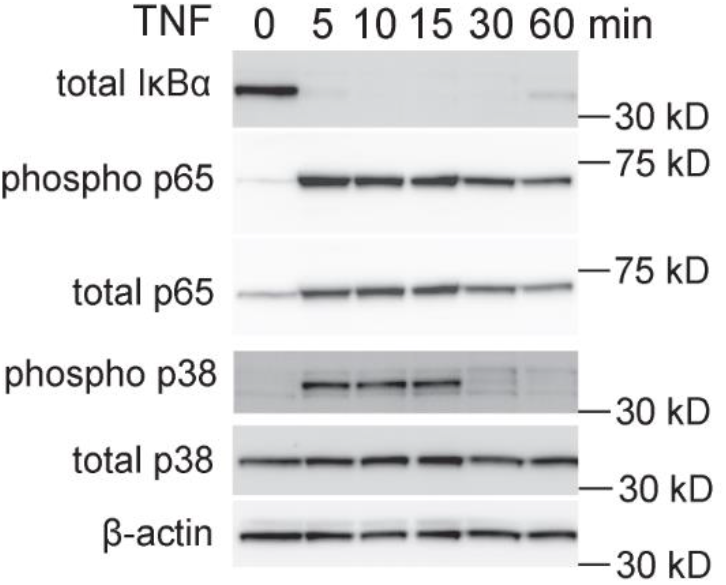
TNF signaling induction. Validation of TNFα signaling induced upon TNF stimulation. U2OS cells were stimulated with 100 ng/μl TNF for the indicated time.

**Supplementary Figure 4.**
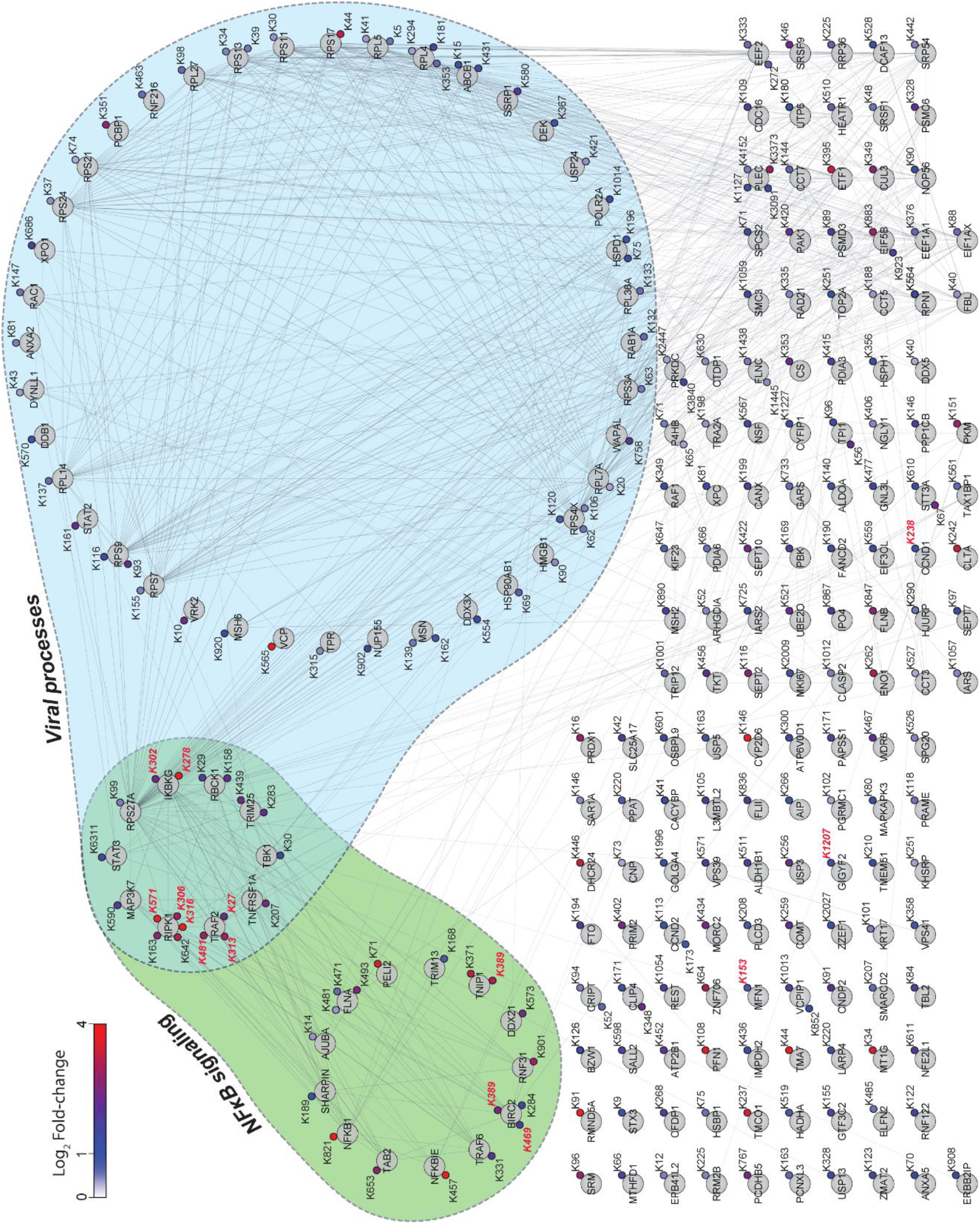
TNF signaling induced ubiquitinome regulation. Cytoscape network of diGly proteins with significantly upregulated diGly sites (DIA; 5% FDR). Green and blue areas mark ubiquitinated proteins associated with NFκB signaling (GO:0043122; GO:0051092) and viral processes (GO:0016032), respectively. Upregulated diGly sites also identified by DDA (5% FDR) are marked in red fond.

**Supplementary Figure 5.**
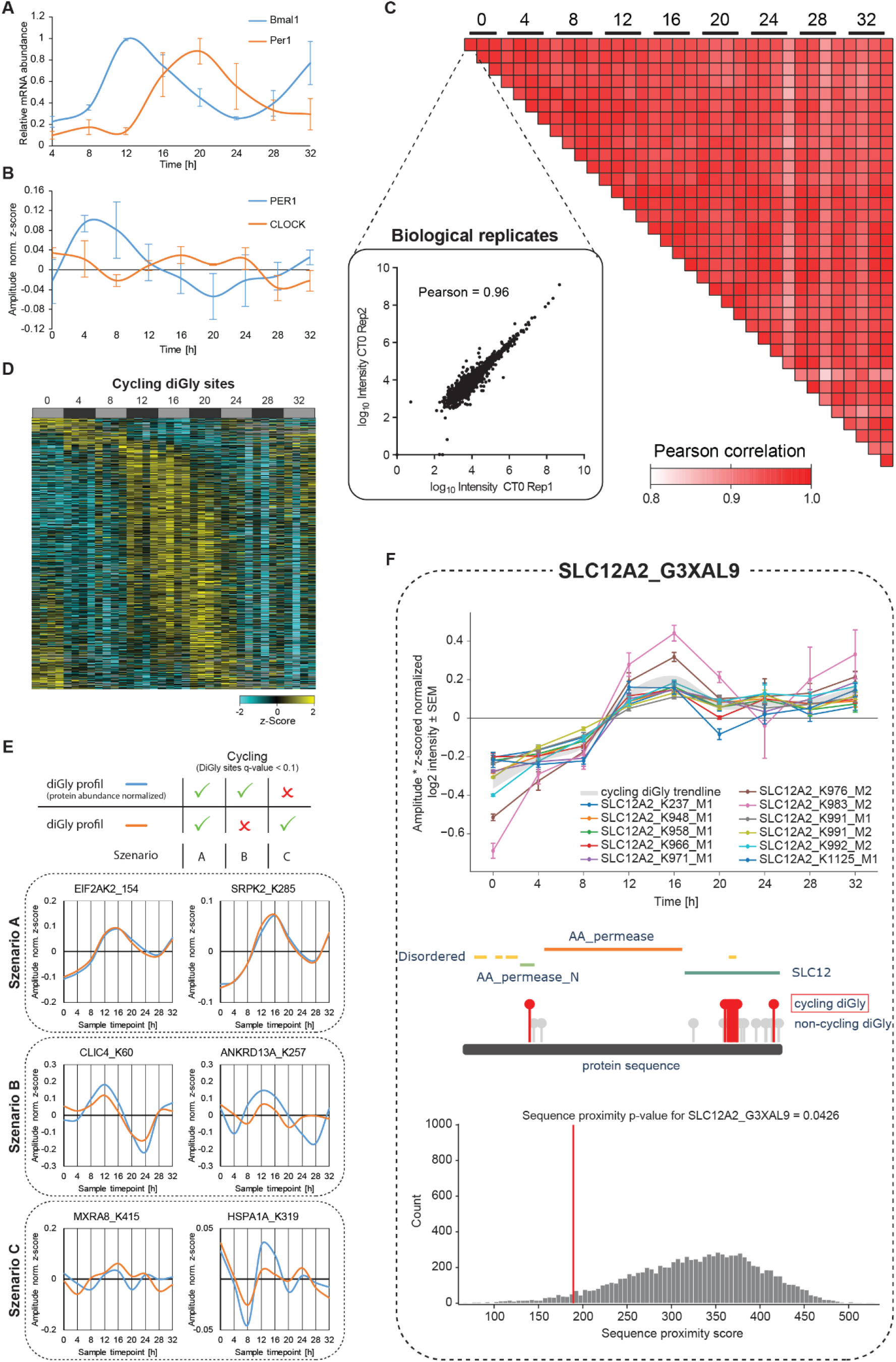
Circadian regulation of the ubiquitinome. **a** Average mRNA abundance (±SEM) of Bmal1 (blue) and Per1 (orange) relative to a Gapdh control in each time point of three biological replicates. **b** Amplitude normalized z-scores of protein profile of core clock proteins Per1 and CLOCK. (C) DiGly proteome correlation matrix (Pearson correlation). Scatter plot shows the correlation of a representative biological replicate. **d** Heat map of intensities (log2 z-score normalized) of ubiquitination sites (row) over time (columns) ordered by phase of oscillation. **e** Various outcomes for protein abundance normalization for diGly sites on diGly site profiles (upper panel). Median, amplitude normalized z-score values for diGly profiles with and without protein abundance normalization are shown in specific example plots (lower panels) with blue and orange lines, respectively. **f** Example of the proximity analysis of cycling ubiquitin clusters (http://cyclingubi.biochem.mpg.de). Cycling sites (q-value < 0.1) (top) and their location in the protein sequence along with its domain annotation (middle) and proximity score (average distance, p-value < 0.1) (bottom) are displayed for SLC12A2.

